# Exploring the Pharmacological Potential of Kaurenoic Acid Produced via Synthetic Biology

**DOI:** 10.1101/2024.10.10.617616

**Authors:** Lígia L. Pimentel, Francisca S. Teixeira, Ana M. S. Soares, Paula T. Costa, Ana Luiza Fontes, Susana S. M. P. Vidigal, Manuela E. Pintado, Luis M. Rodríguez-Alcalá

**Affiliations:** Universidade Católica Portuguesa, CBQF — Centro de Biotecnologia e Química Fina — Laboratório Associado, Escola Superior de Biotecnologia, Rua Diogo Botelho 1327, 4169-005 Porto, Portugal

**Author notes:** Correspondence (L.L.P.); (L.M.R-A.). (L.L.P.); (F.S.T.); (A.M.S.S.); (P.T.C.); (A.L.F.); (S.S.M.P.V.); (M.E.P.); (L.M.R-A.). Those authors contributed equally to this work (Lígia L. Pimentel, Francisca S. Teixeira).

**Keywords:** Kaurenoic acid, QSAR, bioactive compound, diterpene, antimicrobial activity, anti-inflammatory activity

## Abstract

**Background:** Kaurenoic acid (KA) is a bioactive diterpenoid commonly found in traditional medicinal plants such as Copaifera species, widely used in Amazonian ethnomedicine for its anti-inflammatory and antimicrobial properties. However, the sustainable supply of KA is limited due to environmental pressures and its complex extraction process from natural sources. Synthetic biology presents an innovative solution for producing KA, potentially reducing environmental impact while maintaining its traditional medicinal value.

**Aim of the study:** This work discusses the potential bioactive properties of kaurenoic acid (KA) obtained through synthetic biology. While ethnopharmacological studies have highlighted the anti-inflammatory and antimicrobial effects of plant extracts containing KA, limited research has focused on the pure compound due to its cost and limited availability.

**Materials and methods:** The study employed quantitative structure-activity relationship (QSAR) modeling and in vitro assays to investigate the anti-inflammatory and antimicrobial activities of S-KA and KNa. Physicochemical characterization, including Fourier transform infrared spectroscopy (FTIR-ATR), differential scanning calorimetry (DSC), and X-ray diffraction (XRD), was conducted to compare structural properties and purity with G-KA. Additionally, solubility studies were performed across various solvents to assess the potential for different bioapplications.

**Results:** Purity assessments revealed 99.06% for S-KA versus 98.83% for a commercial standard. In silico calculations indicated that KA is hydrophobic. Solubility tests prompted the synthesis of a sodium salt derivative (KNa), increasing water solubility. Experimental evaluations demonstrated that S-KA and KNa exhibited similar (IL-6, IL-8) anti-inflammatory activity compared to betamethasone, and growth inhibition was observed against Staphylococcus aureus and Staphylococcus epidermidis.

**Conclusion:** These findings highlight that kaurenoic acid from fermentation offers a sustainable alternative to naturally sourced KA with comparable bioactivity. The sodium salt derivative (KNa) enhances water solubility, expanding its potential for pharmacological applications. This study highlights the relevance of synthetic biology in preserving traditional medicinal knowledge while promoting environmental sustainability.

## 1. Introduction

Nowadays, one of the health problems to be addressed is the increment of deaths (41 million/year; 77% of total deceases) related to noncommunicable diseases (NCDs; diabetes, cancer, respiratory, and cardiovascular diseases) (World Health Organization, 2018). In general, once manifested, these conditions require medication to be controlled. On the other hand, such drugs can exert side effects and are obtained through chemical synthesis (i.e. high energy demand and pollution risk). Thus, in current pharmacopoeia and medicine, there is a growing demand for natural biomolecules to address such health challenges. They are sought for their potential biocompatibility and biodegradability, offering alternatives to synthetic counterparts. As a result, consumers perceive such compounds as safer and greener options.

To meet this need sustainably, these biomolecules should be produced through environmentally friendly processes. While plant secondary metabolites are excellent candidates, their traditional production methods often increase pressure on agricultural systems, requiring land, water, and fertilizers, thus competing with food production. In response, advances in synthetic biology have enabled the engineering of microorganisms as cell factories for large-scale biomolecule production, aligning with green chemistry principles.

A paradigmatic example of such valuable molecules is Kaurenoic acid. Also known as ent-kaur-16-en-19-oic acid, kauren-19-oic acid, or cunabic acid, is a diterpenoid derived from ent-kaurene and a scaffold for the synthesis of the phytohormones gibberellins. This explains why since the first report of ent-kaurene in the leaves of a New Zealand pine (locally called kauri), more than 1300 derivatives have been described (Ding et al., 2017). These biomolecules have attracted great interest due to their bioactive properties; specifically, kaurenoic acid has been associated with antinociceptive, anti-inflammatory, antiasthmatic, and antimicrobial activities (Zhao et al., 2022). Nonetheless, Amazonia people highly value the resins/pitch from *Protium* species also known as “breu branco” and “breu preto” as well as those from *Copaifera* (i.e. Copaíba resin) for the relief of headaches (da Cruz Albino et al., 2021), against cystitis, skin and mucosa infections (da Trindade et al., 2018). As bioactives, breu pitches contain α- terpineol, α-amyrin, β-amyrin, α-amyrone, and β-amyrone (da Cruz Albino et al., 2021) while copaiba resin is characterized by the presence of β-caryophyllene (up to 55%) and kaurenoic acid (up to 44.3%) depending on the source (da Trindade et al., 2018).

Under these circumstances, kaurenoic acid could be highly valuable in the study and development of new medicines, nutraceuticals, and cosmeceuticals, which are a major priority right now due to the current health and environmental issues. However, exploiting the Amazon’s natural resources for pharmacological purposes will increase the existing environmental pressure on this tropical forest (Gatti et al., 2021). In the case of some resins, such as copaíba, the tree will produce (under the best conditions) 0.5 L/d per year (da Silva Medeiros and Vieira, 2008). It will require further purification by molecular distillation (Galúcio et al., 2022).

Moreover, these kinds of studies also bring the opportunity to conserve the traditional knowledge of indigenous communities who have used these resins for generations. Preserving this knowledge enhances our understanding of these bioactives and their sources, while also underlines the importance of responsible research practices that respect and protect biodiversity.

In the research related to kaurenoic acid, two situations can be observed: either the compound is obtained as a plant extract, a mixture of ent- kaurenes/Gibbellerines, or it is produced through chemical synthesis (Ding et al., 2017). Therefore, this greatly limits the amount available to perform different simultaneous studies. In this situation, synthetic biology, when carried out on a semi-industrial scale, represents a very relevant opportunity since it can offer, in comparison, larger quantities while using processes where it is possible to control the purity of the biomolecule. It also allows the application of sustainable and less environmentally aggressive methods.

Given that synthetic biology enables high-yield and cost-effective production of novel biomolecules—thereby enhancing their availability in terms of purity and quantity—this study aims to assess any potential physical and chemical differences between commercially available kaurenoic acid and that produced via synthetic biology. Additionally, we seek to understand the biocompatibility, anti-inflammatory, and antimicrobial activities of the latter. We hypothesize that synthetic biology-derived kaurenoic acid will not differ chemically from that obtained through other sources and will exhibit biological properties consistent with those reported in the literature. This is the first study to explore the pharmacological potential of synthetic kaurenoic acid, providing a sustainable approach to its large-scale production.

## 2. Materials and Methods

### 2.1. Chemicals and Samples

Within the scope of this work, two samples of kaurenoic acid (KA) were studied: G-KA (1 mg, ≥ 95% purity) was purchased from Glentham Life Sciences (Corsham, United Kingdom) and S-KA, from synthetic biology, was kindly provided by Amyris, Inc. (Emeryville, CA, USA). Furthermore, the sodium salt derived from kaurenoic acid, KNa, was synthesized using *n-*hexane (Hxn) (HPLC Grade, 97%) from VWR Chemicals (Radnor, Pennsylvania, USA) and sodium hydroxide (NaOH) 1 M (aq.) from Fisher Scientific (Pittsburgh, PA, USA).

For solubility studies the solvents used were acetone (AcO) (HPLC Grade ≥ 99.8%) from Fisher Scientific (Pittsburgh, PA, USA), dichloromethane (DCM) (HPLC grade, ≥ 99.9%) from VWR Chemicals (Radnor, Pennsylvania, USA), ethanol (EtOH) (Food grade, 96% v/v) from Panreac Applichem ITW Reagents (Darmstadt, Germany), dimethyl sulfoxide (DMSO) (anhydrous, ≥ 99.9%) and ethyl acetate (EtOAc) (Food grade, ≥ 99%) from Merck (Darmstadt, Germany), and ultra-pure water was obtained through a Milli-Q system (Merck Millipore, Burlington, MA, USA) coupled with a 0.22 μm membrane filter (Millipak; Merck Millipore).

For GC-MS analysis all the samples were derivatized with *N,O*- Bis(trimethylsilyl) trifluoroacetamide with 1% trimethylchlorosilane (BSTFA) purchased from Merck (Darmstadt, Germany).

For LC-ESI-QTOF-MS/MS analysis, all reagents were LC-MS grade, being isopropanol (IPA), acetonitrile (ACN), formic acid, and ammonium formate purchased from VWR (Radnor, PA, USA).

For cytotoxic assays, PrestoBlue assay kit was purchased from Thermofischer (Waltham, Massachusetts, EUA). To perform the anti-inflammatory assays, lipopolysaccharides (LPS) from *Escherichia coli* O111:B4 was obtained from Merck (Darmstadt, Germany), and IL-10, IL-6, IL-8 and TNF-α by Biolegend (San Diego, California, U.S.). For total cellular protein quantification, Pierce™ BCA Protein Assay Kit was purchased from Thermofischer (Waltham, Massachusetts, EUA).

### 2.2. Synthesis of Kaurenoic sodium salt (KNa)

The sodium salt of kaurenoic acid (KNa) was synthesized by dissolution of S- KA (1.07 g, 3.5 mmol) in 100 mL of *n*-hexane. After total dissolution, 50 mL of a 0.5 M NaOH aqueous solution were added, and the mixture was kept under stirring for 1 h at room temperature (Fig S1). KNa precipitated as a white solid in the aqueous phase and was recovered by filtration. The solid was washed with cold water and dried overnight at 60 °C to provide the desired compound.

### 2.3. Quantitative Structure-Activity Relationship (QSAR) predictions

To estimate the potential biological activity of the kaurenoic acid, key properties were taken into consideration namely, the octanol-water partition coefficient (LogP), human skin permeability coefficient (Log Kp), the probabilities of skin sensitization and Ames toxicity, as well as the estimation of its most probable binding target. These properties were consulted in reliable online chemoinformatics that predicts them based on different QSAR models (*in silico* calculations), using the canonical SMILES of kaurenoic acid (CC12CCCC(C1CCC34C2CCC(C3)C(=C)C4)(C)C(=O)O).

Usually, QSAR models predict a series of physicochemical and biological properties of chemicals based on their main physicochemical properties such as molecular weight (MW), topological polar surface area (TPSA), density and water solubility. In this case, the consulted databases and tools were PubChem (https://pubchem.ncbi.nlm.nih.gov), Chemspider (https://www.chemspider.com), OCHEM (https://ochem.eu/home/show.do), SwissADME (Daina et al., 2017), pkCSM (Pires et al., 2015), ADMETlab (Dong et al., 2018), SwissTargetPrediction (Daina et al., 2019), BindingDB (https://www.bindingdb.org/rwd/bind) and SEA (Keiser et al., 2007).

### 2.4. Physicochemical characterization

#### 2.4.1. Solubility of KA and KNa

The solubility of S-KA and KNa molecules was assessed in different solvents such as water, dimethyl sulfoxide (DMSO), a mixture of 1% (v/v) of DMSO in water, ethanol (EtOH), ethyl acetate (EtOAc), acetone (AcO), hexane (Hxn) and dichloromethane (DCM). Accurately weighed amounts of S-KA or KNa were gradually added to 1 mL of each solvent; the mixture was vortexed after each addition until solid precipitation was noticed. Then, the solution was collected, and the precipitate was dried and weighed. The solubility was calculated by the difference between the mass of the compound added and the mass of the compound that did not dissolve, in 1 mL of solvent. Tests were performed in duplicate.

#### 2.4.2. Fourier Transform Infrared Spectroscopy-Attenuated Total Reflectance (FTIR-ATR)

The FTIR-ATR analysis of the kaurenoic acid and sodium salt samples was performed on a Perkin Elmer Paragon 1000 FTIR (Waltham, Massachusetts, United States) with the ATR accessory. The spectra were obtained in the wavenumber range of 4000-550 cm^-1^, with a resolution of 4 cm^-1^, by accumulating 16 scans. The FTIR-ATR vibrational bands were identified based on literature (Socrates, 2001) and are summarized in Table S1.

#### 2.4.3. Differential Scanning Calorimetry (DSC)

Measurements of DSC were performed using a NETZSCH DSC 204 F1 Phoenix (NETZSCH-Gerätebau GmbH, Selb, Germany) calorimeter. The samples (2-4 mg) were prepared in duplicate by weighing them into aluminum crucibles and sealing it. Runs were performed by heating each sample, in duplicate, from -20 °C to 500 °C, with a heating rate of 10 °C/min and including an isothermal step at 500 °C for 1 min at the end of the run. A nitrogen flow rate of 100 mL/min was kept during the DSC runs. An empty and sealed crucible was used as a reference.

#### 2.4.4. X-ray powder diffraction (XRD)

X-ray experiments were performed on a MiniFlex 600 diffractometer (Rigaku Europe SE, Neu-Isenburg, Germany) supplied with Cu-Kα radiation (λ = 0.15418 nm, 40 kV, 15 mA). Samples were scanned in duplicate from 3° to 90° (2*θ*) in steps of 0.01° at a speed rate of 3°/min.

### 2.5. Liquid chromatography electrospray ionization quadrupole time-of-flight (LC-ESI-QTOF)

The samples of S-KA and G-KA were dissolved in IPA:ACN (9:1, v/v) at 0.25 mg/mL and analyzed on an UHPLC instrument (Elute; Bruker, Billerica, MA, USA), equipped with an Acquity UPLC BEH C18 (17 µm) pre-column (Waters, Milford, MA, USA), an Intensity Solo 2 C18 (100 x 2.1 mm) column (Bruker), and coupled with an UHR–QTOF detector (Impact II; Bruker). The injection method was based on conditions reported by Sarafian et al (2014) and Calderon et al. (2019), with some modifications previously described by Teixeira et al., (2023). Thus, mobile phases consisted of ACN:upH_2_O (6:4, v/v) (Phase A) and IPA:ACN (9:1, v/v) (Phase B), each one added with 0.1% (v/v) formic acid and 10 mM ammonium formate. Phase B gradient flow was set as follows: 0.0 min: 40%, 2.0 min: 43%, 2.1 min: 50%, 12.0 min: 54%, 12.1 min: 70%, 18.0 min: 99%, 20.0 min: 99%, 20.1 min: 40% and 22 min: 40%. Flow rate was set at 0.4 mL/min and column temperature at 55 °C. The injection volume was 3 µL in positive ionization mode and 5 µL in negative ionization mode. For MS analysis, the following parameters were applied: end plate offset voltage 500 V, capillary voltage 4500 V (positive ionization) or 3000 V (negative ionization), nebulizing gas pressure 35 psi, drying gas flow 8 L/min, drying gas temperature 325 °C, quadrupole ion energy 3 eV (positive ionization) or 5 eV (negative ionization), Collision energy 10 eV (positive ionization) or 5 eV (negative ionization). Acquisition was performed in an auto MS/MS scan mode over a mass range of m/z 50-1500. For both ionization modes, an external mass calibration was performed with a solution of IPA:upH_2_O (1:1, v/v) added with 0.2% (v/v) formic acid and 0.6% (v/v) NaOH 1M, continuously injected at 180 µL/h.

#### 2.5.1. Gas chromatography-mass spectrometry (GC-MS)

The samples (S-KA, G-KA and KNa) were derivatized into their trimethylsilyl derivatives by accurately weighing 1 mg of sample and adding 500 uL of DCM and 30 uL of BSTFA. After incubation at 30 °C for 60 minutes, DCM was added to a final volume of 1.5 mL. The derivatized samples were analyzed on a GC-MS model EVOQ (Bruker, Karlsruhe, Germany) coupled to a mass spectrometer, with a Rxi- 5Sil MS column (30m × 250 µm × 0.25 µm) at constant flow of 1 mL/min. The carrier gas used was helium and the GC-MS conditions were as described by Teixeira *et al*. (2021). The purity of the samples was assessed by area percent.

### 2.6. Cell culture

The human monocytic cell line THP-1 (ATCC [TIB-202]) was kept in culture in RPMI-1640 Media (Gibco) supplemented with 10% Fetal Bovine Serum (FBS) (Gibco), 1% penicillin-streptomycin antibiotic (Gibco) and 50 mM of beta-mercaptoethanol (Gibco) at 37 °C, with 5% CO2 and humified atmosphere. For the experiments, THP-1 cells were seeded and differentiated into macrophages by treatment with 50 nM of Phorbol 12-myristate 13-acetate (PMA) (Sigma) for 48 h.

### 2.7. Cytotoxicity assays

Cytotoxicity of S-KA and KNa on macrophages was evaluated using PrestoBlue assay according to the manufacturer’s instructions. THP-1 cells were seeded at 1x10^4^ cells/well in 96-well plates (Thermofischer, Waltham, Massachusetts, EUA) and differentiated into macrophages. Cells were exposed to samples at different concentrations (0.1 to 1 mg/mL in medium containing 1% DMSO) for 24 h, in quadruplicates. Wells without cells and containing S-Ka and KNa were used to subtract a possible influence of these samples in the PrestoBlue fluorescence signal. Cells treated with 10% of DMSO were used as a negative control. After incubation, PrestoBlue was added to the media and incubated for 2 h. The fluorescence signal was read in a Synergy H1 microplate reader (BioTek, Winooski, Vermont, EUA). Results were expressed as the percentage of metabolic inhibition compared to the control (cells without treatment). Two independent experiments were performed. The human keratinocyte cell line HaCaT (CLS - Cell Line Services - 300493) was kept in culture in Dulbecco’s Modified Eagle Medium (DMEM) supplemented with 10% FBS and 1% penicillin-streptomycin antibiotic at 37 °C, with 5% CO_2_ in a humidified atmosphere. Cytotoxicity of samples on human immortal keratinocytes (HaCaT) was also evaluated using the abovementioned protocol, with some modifications. Cells were seeded at 1x10^4^ cells/well in 96-well plates and exposed to the samples at different concentrations (0.13 to 0.01 mg/mL) diluted in DMEM for 24 h, in quadruplicates. Wells with media supplemented with the samples (without cells) were used to subtract a possible influence of the samples in the PrestoBlue fluorescence signal. Cells treated with 10% DMSO were used as a negative control. Two independent experiments were performed.

#### 2.7.1. Anti-inflammatory activity by Enzyme-Linked Immunosorbent Assay (ELISA)

As previously described by Teixeira *et al*. (2021) with few modifications, THP-1 cells seeded at 3x10^5^ cells/well in 24-well plates (Thermofischer, Waltham, Massachusetts, EUA), were differentiated into macrophages. Cells were treated for 24 h with S-KA and KNa at 60 μg/mL in the presence or absence of lipopolysaccharides (LPS) from *Escherichia coli* O111:B4 (Merck, Darmstadt, Germany) at 0.1 µg/mL to induce inflammation. For anti-inflammatory control, cells were treated with 20 μM of betamethasone. Medium supernatants were collected and used to evaluate the levels of pro-inflammatory cytokines human IL-6, IL-8, TNF-α (Tumor Necrosis Factor Alpha) and anti-inflammatory IL-10 by ELISA (Biolegend, San Diego, California, U.S.). To normalize cytokine levels, total cell protein was determined. Cells were lysed with water and used for protein quantification via Pierce™ BCA Protein Assay Kit (Thermofischer, Waltham, Massachusetts, EUA). Results were reported as percentage of cytokine expression. For all measurements, the LPS-induced inflammation values (in pg Interleukin/μg cell protein) on macrophages were considered as the maximum expression (100%). Therefore, the percentage of interleukin expression for S-KA, KNa and betamethasone exposure was normalized considering the maximum expression by LPS. Two independent experiments were performed.

### 2.8. Antimicrobial activity evaluation

#### 2.8.1. Microorganisms

The antimicrobial activity of kaurenoic acid (S-KA) and kaurenoic sodium salt (KNa) was determined following the agar microdilution method described by Golus *et al*. (2016). The reference strains tested were *Escherichia coli* (DSM 1576), *Pseudomonas aeruginosa* (DSM 1128), *Staphylococcus aureus* (DSM 799) and S*taphylococcus epidermidis* (LMG 10474) to determine the antibacterial activity and the yeast *Candida albicans* (DSM 1386) to assess the antifungal activity of the samples.

The inocula were prepared according to the guidelines of the Clinical Laboratory Standards Institute (CLSI-M07-A9 standard). Thus, isolated colonies from each microorganism were suspended in Müeller-Hinton broth (MHB, Biokar Diagnostics, Beauvais, France) to achieve a turbidity equivalent to 0.5 MacFarland standard (∼1x10^8^ CFU/mL, Optical density ranging from 0.08-0.1 at 625 nm). These microbial suspensions were diluted 1:10 (v/v) in MHB to obtain a concentration of 10^7^ CFU/mL.

#### 2.8.2. Samples

Stock solutions of S-KA (25 mg/mL) and for KNa (12 mg/mL) were prepared in DMSO. Dilutions of these samples were prepared in Müeller-Hinton agar (MHA, Biokar Diagnostics, Beauvais, France) to reach different concentrations. S-KA and KNa tested concentrations ranged from 0.0062-0.4 mg/mL for *S*. *aureus* and *S*. *epidermidis*. For *E. coli*, *P. aeruginosa* and *C*. *albicans*, tested concentrations for KA ranged from 0.08-1.25 mg/mL and for KNa from 0.08-0.65 mg/mL. A sterility control (negative control only with culture medium MHA), a growth control (MHA with inoculum for each microorganism) and DMSO controls [concentrations reached maximum 5% (v/v)] were also included. Phenoxyethanol (PE), a preservative used in many cosmetic products, was prepared in MHB (stock solution at 28 mg/mL) and used as positive control; concentrations tested ranged from 0.325-12 mg/mL. The Eppendorf centrifuge tubes were vortexed and kept at 50°C in a block heater (Stuart SBH2000D/3, Cole-Parmer Ltd, UK) until the samples were transferred to the microplate.

#### 2.8.3. Procedure

From each sample concentration to be tested, 100 µL were pipetted in triplicate into a 96-well microplate with round base (Sarstedt, Nümbrecht, Germany). Sterility, growth, DMSO and PE controls were also pipetted in triplicate into the microplate. After the agar solidification, 2 μL of each inoculum prepared at 10^7^ CFU/mL were applied to the agar of the respective wells of the microplate. The plates containing bacterial strains were incubated at 37 °C for 16-18 h and those with the yeast *C. albicans* were incubated at 30 °C for 24-48 h. The MIC was registered as the lowest concentration that completely inhibits the visible microorganism growth.

#### 2.8.4. Statistics

Results are reported as mean values ± standard deviation. Data were first analysed for normality distribution (i.e., Shapiro-wilk). Levene’s test was applied to verify the homogeneity of the variances. Afterwards, one-way ANOVA test was applied with Tukey post hoc test to determine differences within groups. Level of significance was set at 0.05. Analyses were performed with the aid of the IBM SPSS Statistics software (28.0 version, Chicago, USA).

## 3. Results and Discussion

### 3.1. Quantitative Structure-Activity Relationship (QSAR)

To estimate the potential biological activity of the kaurenoic acid, key properties were consulted in reliable online chemoinformatics tools based on different QSAR models, and the results are summarized in Table S2 (please see Supplementary data) and described below.

Assuming the possibility of a topical application for kaurenoic acid, since those are common utilizations of kaurenoic-containing oils, crucial properties were estimated, namely the octanol-water partition coefficient (LogP), the human skin permeability coefficient (Log Kp), and the skin sensitization probability.

Nowadays, a simple generally accepted approach to predict the drug-likeness of a molecule is the well-known “rule of 5” by Lipinsky *et al*. (Lipinski et al., 2001), which states that poor absorption or permeation is more likely when: (1) the MW is over 500 g/mol; (2) the LogP is over 5; (3) there are more than 5 H-bond donors (expressed as the sum of OHs and NHs) and (4) there are more than 10 H-bond acceptors (expressed as the sum of Ns and Os). Considering these four rules and the results presented in Table S2 **Error! Reference source not found.**, kaurenoic acid follows all the parameters to be considered a good candidate compound, except for LogP which raises some doubts depending on the consulted database: SwissADME states a LogP value of 4.47 (under 5, so suitable for biological applications) while Chemspider reports a LogP value of 6.37 (over 5, so not suitable for biological applications). Despite the different LogP values reported, the fact that both were positive indicates that kaurenoic acid is a hydrophobic molecule. In fact, the same happens to a known commercial cosmetic ingredient, squalene, that also respects the Lipinski “rule of 5” except for LogP value that varies from 8.62 (OCHEM) to 13.09 (Chemspider). This suggests that a single deviation from the Lipinsky “rule of 5” is not an impediment to a molecule show a good physicochemical performance.

Regarding the log Kp values, different results were obtained depending on the source: -4.29 from SwissADME and -2.735 from pkCSM. Even though the values were different, both revealed to be negative, which indicates good human skin permeability for KA.

Concerning skin sensitizing, most QSAR sources evaluate this property as “Yes or No”, but ADMETlab takes a further step and estimates its probability. For KA, this *in silico* tool, estimated a not significant probability (0.03) for skin sensitization, therefore considering it as a suitable molecule for topical application. Furthermore, ADMETlab also assesses the potential carcinogenic effect of chemicals by the Ames toxicity test, using the bacterial strain *Salmonella typhimurium*, reporting it as a probability. About KA, the calculated probability was 0.016, suggesting that this molecule is not mutagenic.

Moreover, QSAR models such as SwissTargetPrediction, BindingDB and SEA, suggested that KA could be a ligand of corticosteroid 11-beta-dehydrogenase isozyme 1 (11β-HSD1). Besides being involved in insulin signalling, this enzyme regulates cortisol metabolism and is expressed in tissues such as placenta, skin epidermis, liver and adipose tissue (Tomlinson and Stewart, 2001). The regulation of glucocorticoids by the catalysation of the intracellular conversion of cortisone to cortisol by this enzyme is described as its main function, which is directly associated with anti-inflammatory regulation and immunosuppressive processes (Peng et al., 2016).

### 3.2. Physicochemical characterization

#### 3.2.1. S-KA and KNa Solubility

The solubility of a biomolecule plays an important role in its bioactivity since it will determine from the way of administration to absorption and distribution. As commented in the above section, kaurenoic acid is a hydrophobic molecule (i.e. estimated LogP 4.47-6.37, Table S2). In an attempt to obtain a form of kaurenoic acid more soluble in water, the synthesis of the corresponding sodium salt was assayed. The reaction synthesis yield was 93.3%.

The solubility study was assessed in several solvents such as water, DMSO, 1% (v/v) DMSO in water, EtOH, EtOAc, AcO, Hxn and DCM, to evaluate the solubility of S-KA and KNa for bioapplications and/or to evaluate their solubility for organic synthesis purposes. The results are summarized in Table 1, showing that KNa, as expected, was more soluble in water than S-KA (3.870 ± 0.060 mg/mL *vs* 0.013 ± 0.003 mg/mL, respectively). Since DMSO is a common solvent used for *in vitro* tests, it was also assayed. Thus, it was observed the opposite behaviour: S-KA revealed higher solubility than KNa (357.345 ± 1.535 mg/mL *vs* 12.730 ± 0.280 mg/mL, respectively).

**Table 1.**
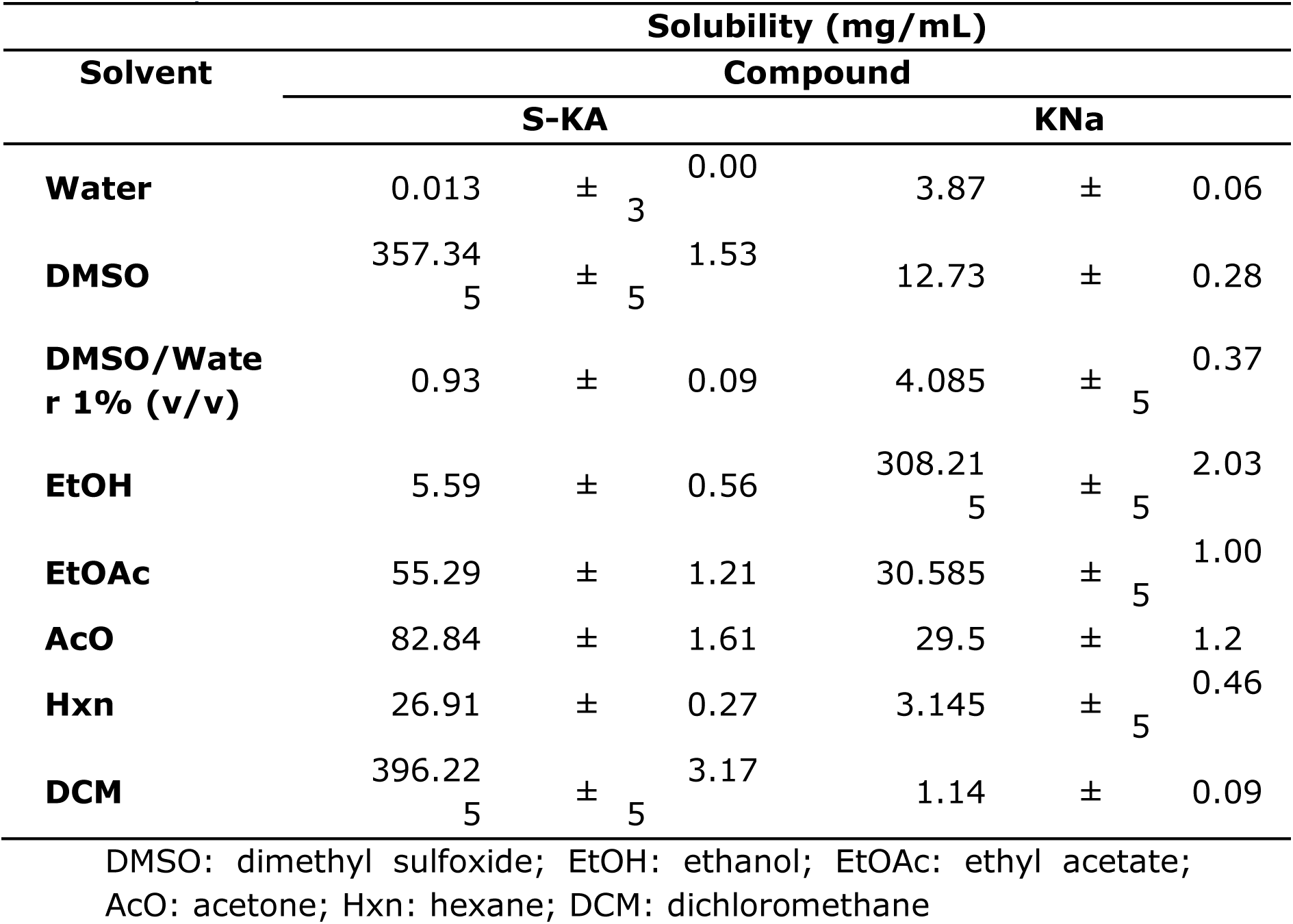
Solubility of S-KA and KNa.

Because of toxicity limitations, in cell lines studies it is common not to exceed 1% DMSO in aqueous solutions and this condition was therefore studied. The solubility results for S-KA and KNa were 0.930 ± 0.090 mg/mL and 4.085 ± 0.375 mg/mL, respectively.

Regarding the results obtained for the rest of the solvents evaluated: in ethanol (polar and protic solvent) it was possible to observe that KNa was much more soluble than S-KA (308.215 ± 2.035 mg/mL and 5.590 ± 0.560 mg/mL, respectively). In the other organic solvents studied it was found the opposite behaviour, being the solubility of S-KA always higher than that of KNa. In polar and aprotic solvents such as EtOAc the solubility of S-KA and KNa was 55.290 ± 1.210 mg/mL and 30.585 ± 1.005 mg/mL, and in AcO of 82.840 ± 1.610 mg/mL and 29.500 ± 1.200 mg/mL, respectively. In apolar solvents such as Hxn, the solubility values were 26.910 ± 0.270 mg/mL to S-KA and 3.145 ± 0.465 mg/mL to KNa. In the case of a chlorinated solvent such as DCM, the solubility of S-KA and KNa was 396.225 ± 3.175 mg/mL and 1.140 ± 0.090 mg/mL, respectively.

The obtained results showed that KNa could be a viable alternative to S-KA and helpful for the study of the bioactivity of this molecule since its solubility in water increases (∼300x).

#### 3.2.2. FTIR-ATR analysis

The FTIR-ATR spectra of the synthetic KA (S-KA) and the commercial standard G-KA (natural extract) were compared (Fig 1) to understand if there was any structural difference between them due to their distinct origin.

**Fig 1.**
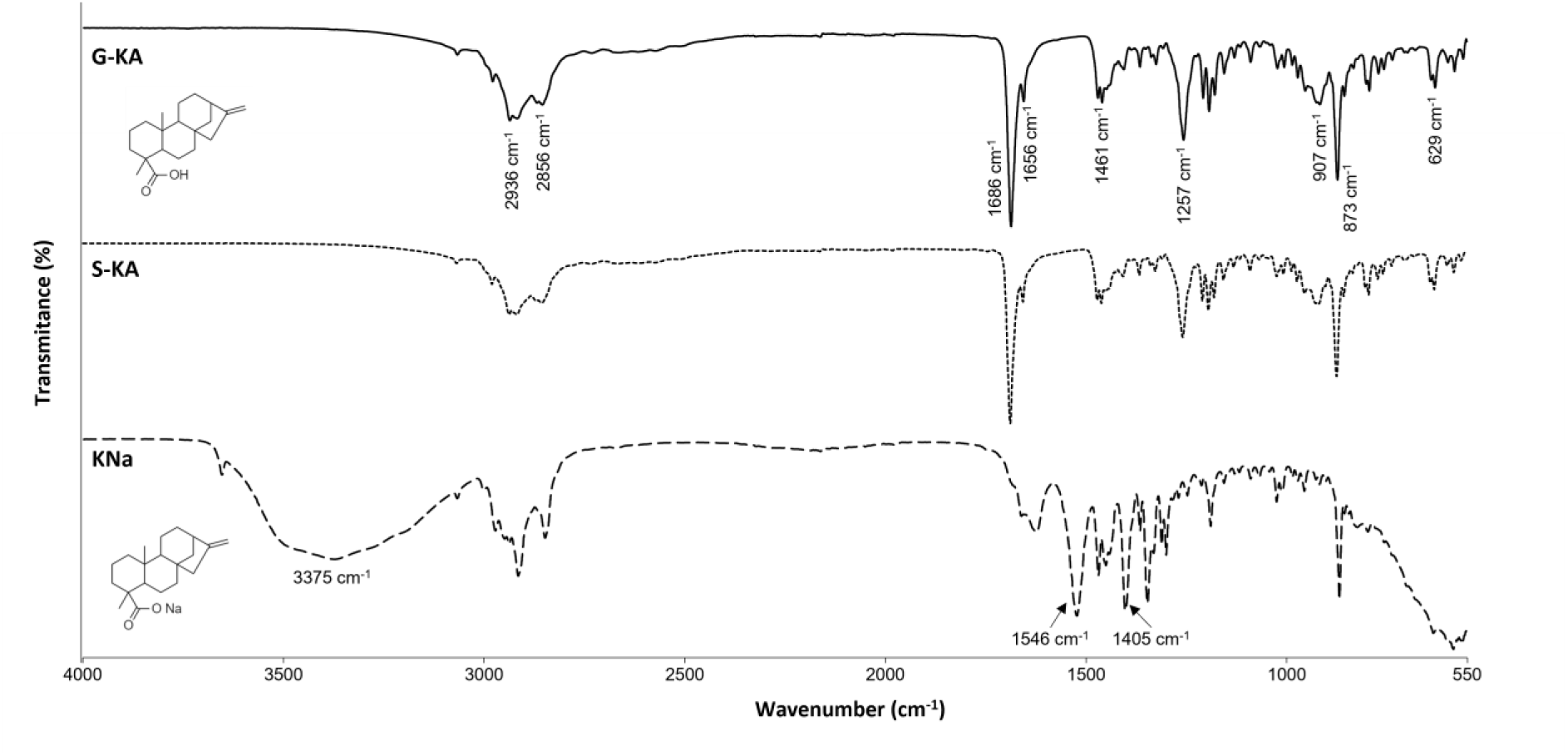
Overlay of the FTIR-ATR spectra of G-KA (solid line); S-KA (dotted line) and KNa (dashed line). Inset: Structures of the kaurenoic acid (upper left) and its sodium salt (lower left).

The FTIR-ATR spectra of the S-KA and the G-KA samples are completely superimposable, evidencing their structural similarity despite their different source. As shown by the structures presented in Fig 1 **Error! Reference source not found.**, the kaurenoic acid molecule is characterized by a cyclic structure with saturated and unsaturated carbon bonds, being the main functional group of carboxylic acid. Ahead, the obtained FTIR vibrational bands are described, confirming that the studied molecules (S-KA and G-KA) present the described structure.

The presence of the FTIR-ATR vibrational bands at 1686 cm^-1^ (C=O stretching), at 1257 cm^-1^ (C-O stretching), at 873 cm^-1^ (O-H out of plane deformation) and at 629 cm^-1^ (C=O deformation), associated to the carboxylic acid group, highlights that the analysed samples correspond to carboxylic acids. The band at 1656 cm^-1^ associated with the C=C stretching vibration, confirms the presence of unsaturated bonds in the molecule. Furthermore, the wider bands, from 2936 cm^-1^ to 2856 cm^-1^ and at 1461 cm^-1^ correspond respectively, to the – CH_2_ stretching and deformation vibrations on aliphatic chains, and finally, the band at 907 cm^-1^ (C-C stretching) related to cyclic aliphatic chains, confirms their prevalent cyclic structure.

As described in the methods section, KNa was synthesized from the S-KA. Comparing the FTIR-ATR spectra of both samples in Fig 1, important structural differences are noticed. Primarily, the appearance of the bands at 1546 cm^-1^ and at 1405 cm^-1^, characteristic of the asymmetric and symmetric vibrations of carboxylate group (COO^-^), respectively, demonstrates the successful synthesis of the salt. Furthermore, the vibrational band at 1686 cm^-1^ (C=O stretching in carboxylic acid groups) shifts to lower wavenumbers and decreases its intensity in the KNa spectrum when compared to the acid; this seems to indicate that the C=O bond of the carboxylic group loses vibrational freedom, since a substantially larger one, Na, replaces the H atom, thus again suggesting the success of the synthesis. Finally, the wide band at 3375 cm^-1^, associated with the O-H stretching vibration in water molecules, is probably due to the presence of residual water within the salt powder, from the synthesis in an aqueous medium. The obtained results were corroborated by Gómez-Hurtado *et al*. (2017) that in the scope of their work synthesized the sodium salt of kaurenoic acid and confirmed both kaurenoic acid and salt structures by NMR and FTIR.

#### 3.2.3. DSC analysis

All the samples in study were subjected to differential scanning calorimetry to evaluate the differences between S-KA, kaurenoic acid obtained by a synthetic procedure, and the commercial standard G-KA. Additionally, the kaurenoic sodium salt (KNa) obtained from S-KA was evaluated. The analysis was performed in duplicate, and the results are presented in Table 2. For both synthesized and commercial kaurenoic acids (S-KA and G-KA, respectively) it was possible to observe the same type of thermal transitions, namely: glass transition at 58.9 ± 11.1 °C and 54.1 ± 0.6 °C, melting transition at 176.5 ± 0.2 °C and 178.6 ± 0.1 °C, and decomposition at 342.2 ± 0.8 °C and 339.6 ± 6.5 °C, for S-KA and G-KA respectively. Regarding these results it is possible to assume that both samples are the same compound, and the melting transition agrees with the literature (Ngamrojnavanich et al., 2003), despite some variations that would be associated to the presence of other kaurene species in the samples. FTIR-ATR results also corroborate these results. For kaurenoic sodium salt (KNa) two melting temperatures at 74.9 ± 1.2 °C and 90.7 ± 7.6 °C and two decomposition temperatures at 438.3 ± 0.6 °C and 479.9 ± 0.4 °C were observed.

**Table 2.**
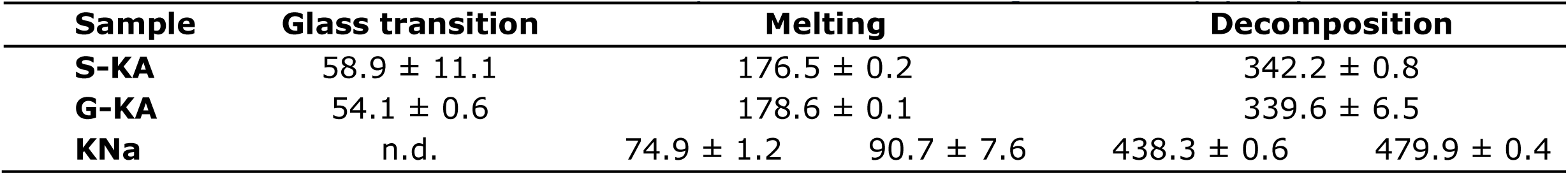
Thermal transitions obtained by Differential Scanning Calorimetry (DSC).

#### 3.2.4. XRD analysis

The X-ray analysis of S-KA and G-KA was performed to evaluate if the physical structure of the kaurenoic acid samples is similar despite the distinct synthesis pathway. This technique is a useful tool to compare the samples of kaurenoic acid (S-KA and G-KA) in terms of purity and crystallinity, by the comparison of the obtained diffractograms. The analysis of the referred diffractograms presented in Fig S2 revealed that both samples (S-KA and G-KA) present high crystallinity due to the appearance of very narrow peaks on the diffractogram. However, and despite having very similar diffractograms, some differences were visible in the intensity of specific peaks. In the case of S-KA (green) a peak at 5° is observed with major intensity than in G-KA. In turn, observing the diffractogram of G-KA (blue) an intense peak at 21° is observed, that is not observed with the same intensity in S-KA. These slight differences could be explained by the presence of other kaurene species in the S-KA, even if in low amounts, that influence the behaviour when exposed to X-ray radiation, resulting in differences in the diffractograms. This behaviour could be corroborated by the information obtained throughout the analysis by GC-MS, where other derivatives of kaurenoic acid were identified (Section 3.4). In the case of kaurenoic sodium salt (KNa, red) obtained from S-KA, the diffractogram present a different behaviour related to the structure of the salt. The peak angles observed show less intensity and in minor number, indicating that we are in the presence of a different structure, resulting in a different diffraction pattern when subject to X-ray irradiation. This result can also corroborate the success of the synthesis procedure to obtain the salt derivative since the differences between the S-KA and KNa diffractograms are perfectly visible and expected.

### 3.3. LC-ESI-QTOF analysis

To confirm the identity of kaurenoic acid in the different samples, a High Resolution-Mass Spectrometry (HRMS) analysis was performed on S-KA and compared with the commercial reference (G-KA). When working in negative mode, it was observed a major ion at *m/z* 301.2166 and a minor one at m/z 625.4193 in S-KA (Fig 2). The major ion corresponds to the deprotonated ion [M-H]^-^ of kaurenoic acid (M_mi_ = 302.2246 g/mol; Table S2) and its corresponding MS/MS showed no fragmentation (Fig 2). On the other hand, MS and MS/MS spectra of G-KA in negative ionization were similar to those of S-KA and also showed no further fragmentation (Fig 2).

**Fig 2.**
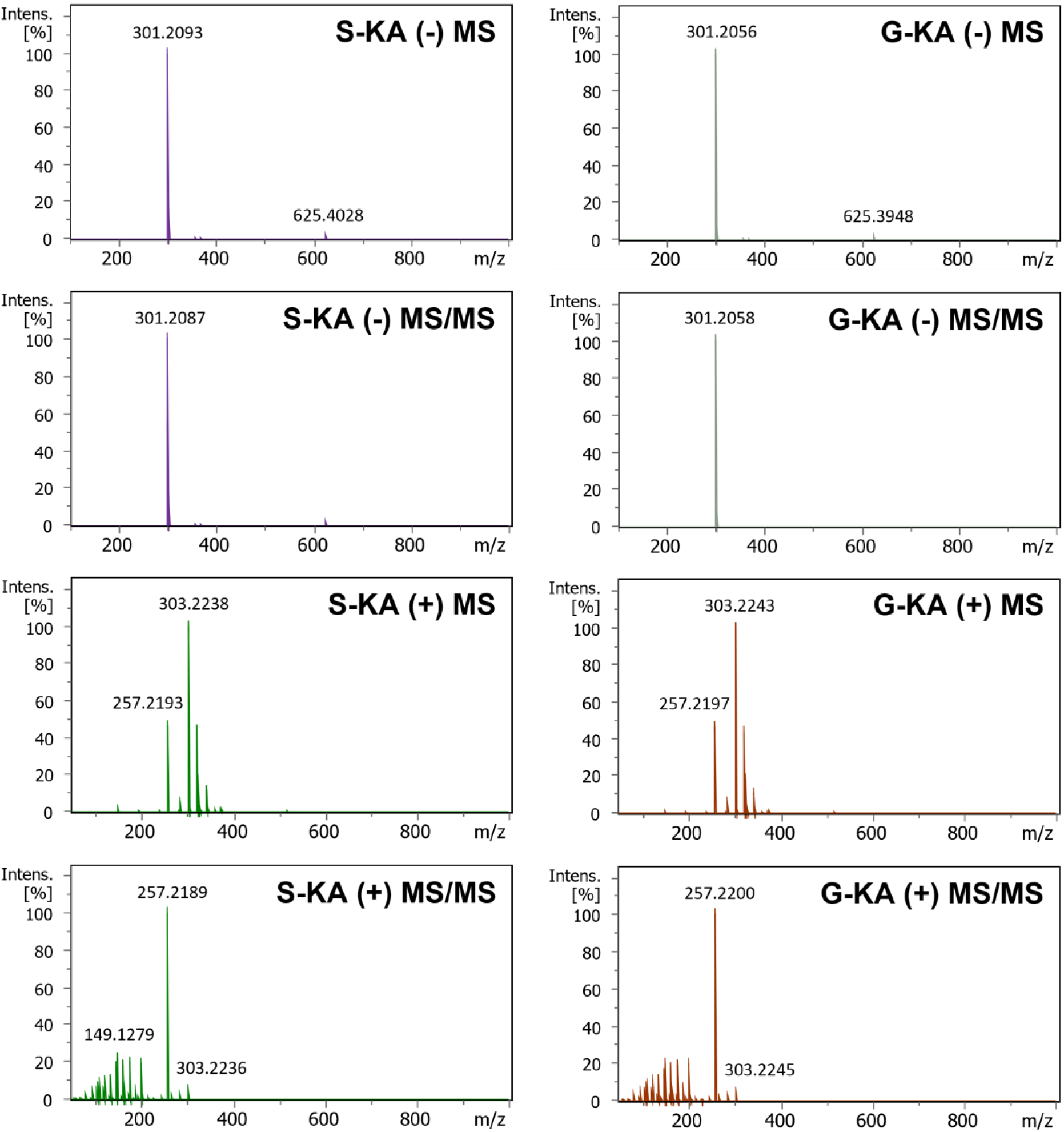
MS and MS/MS spectra of Kaurenoic acid ion obtained by LC-ESI-QTOF-MS/MS in negative ionization mode (-) and positive ionization mode (+) of S-KA and G-KA samples.

Spectra prediction for negative mode run on CFM-ID (data available in https://cfmid.wishartlab.com/queries/65f8dd0161187da1ff343eb942a279599b0 12559) (Wang et al., 2021) showed no MS/MS fragmentation with energies below 10V.

Other authors, when monitoring parent ion at *m/z* 301.2-303.3 in negative mode, but using a triple quad detector, observed no fragmentation of kaurenoic acid at up to 50 eV in MS/MS (Jiang et al., 2019; Miyazaki et al., 2015). Moreover, when both parent and daughter ions were monitored in the last-mentioned research works, the daughter ion of kaurenoic acid at *m/z* 301.2-303.3 was still detected at up to 50 eV. The need to use high energies for MS/MS seems to be a particular feature of kaurenoic acid.

Regarding MS spectra in positive ionization mode, S-KA revealed a major ion at *m/z* 303.2327, being further detected ions at *m/z* 257.2271 and 320.2593 (Fig 2). The MS/MS of ion 303.2327 resulted in an ion at *m/z* 257.267, being also detected minor ions at *m/z* 149.1328 and 303.2326 (Fig 2). Comparing S-KA with G-KA, MS and MS/MS spectra in positive ionization mode were again similar (Fig 2). According to Gasparetto *et al* (2011), the negative ion mode turns out to be the most efficient mode for ionizing kaurenoic acid, however, the obtained results demonstrate that, through the assayed method, it is also possible to analyse kaurenoic acid in positive ionization mode. Moreover, when a spectrum prediction was run on CFM-ID (data available at https://cfmid.wishartlab.com/queries/30d21be85b9130e17c1376685128843c9f2 87e0a) (Wang et al., 2021), the MS/MS spectrum of kaurenoic acid at low energy (10 V) revealed ions at *m/z* 303.2318, 257.2264 and 147.1168, which resemble the ones detected.

### 3.4. GC-MS analysis

The purity of kaurenoic acid in the samples was assessed by GC-MS. According to the obtained data (Table S3), kaurenoic acid was detected as the main compound in all the tested samples as a peak eluting at minute 37.3 min. Since the compound is analysed as a TMS derivative, the mass spectra showed a fragment of 73 *m/z* as the base ion and another fragment of 374 *m/z* (kaurenoic acid MW is 302.5 g/mol) as the molecular ion in all three samples.

In the S-KA sample, purity was 99.06 ± 0.01%, while for G-KA was 98.83 ± 0.04% and it was found to be 99.88 ± 0.01% for KNa. Moreover, besides the major compound, it was possible to identify also ent-kaurene (0.51 ± 0.01%) and ent-kaurenal (0.23 ± 0.01%), in S-KA. When considering G-KA, Kaur-16-en-18- ol, (4α)- (0.27 ± 0.01%) and Kauran-16-ol, (16α)- (0.28 ± 0.01%), were also detected. These are compounds belonging to the ent-kaurene and gibberellin family (Ding et al., 2017; He et al., 2020). In this sample, a peak eluting at minute 35.5 min and accounting 0.45 ± 0.02% was also detected. The mass spectra showed an ion of 73 *m/z* as base ion (therefore suggesting the presence of the hydroxyl group in the molecule) and 388 *m/z* as molecular ion. The NIST database suggested communic acid as the most probable match. However, although the reference spectra from NIST for communic acid (as TMS derivative) displays a fragment of 73 *m/z* as base ion and the molecular one must be 374 (since MW of this molecule is 302.5 g/mol) and accordingly the unknown compound has higher *m/z* value. As G-KA is obtained naturally, it can result from various types of sources such as Copaifera plants, where more than 250 compounds have been described and the terpene fraction is well known (Arruda et al., 2019). Aside from ent-kaurene derivatives, the Copaiba genus produces labdane-type diterpenes where polyalthic and hardwickiic acids are two of the main ones. Both meet the characteristics of the unknown compound: they have the carboxyl group and therefore present a 73 *m/z* ion, and the molecular weight is 316.4 g/mol, thus the molecular ion as TMS derivative would be 388 *m/z*. When comparing the obtained spectra for the unknown compound with the ones from polyalthic and hardwickiic acids, with a computational predictive tool CFM-ID v3 (Allen et al., 2016), a match of 97.0 and 97.5% was observed for the putative compounds, respectively. Finally, no other compound besides ent-kaurenoic acid was identified for KNa; once the sample is derivatized by TMS, the acid form is the one identified in the analysis by GC-MS.

### 3.5. Bioactivity assessment

#### 3.5.1. Cytotoxicity assays

Metabolic Inhibition (%) of KNa and S-KA on macrophages was evaluated in the range of 0.1 to 1 mg/L in RPMI. Differences above 30% of metabolic inhibition were observed for KNa in the concentration value of 0.6 mg/mL that may be inherent to the increase in toxicity (Fig S3). This increased toxicity can be characteristic of the improved water-solubility of KNa, when compared to S-KA. Hence, as the application under study infers the dermal use, the non-toxic concentration in human keratinocytes cell line HaCaT (CLS - Cell Line Services - 300493) was evaluated. The non-toxic concentration determined for both S-KA and KNa was 60 μg/mL (Fig S4). Thus, for the anti-inflammatory assays, the tested concentration was set at 60 μg/mL for both compounds.

#### 3.5.2. Anti-inflammatory activity

The results obtained for the anti-inflammatory response of the tested samples regarding cytokine (human IL-8, IL-6, TNF-α and IL-10) expression are shown in Fig 3. The cytokine inhibitory activity (%) and cytokine concentration (pg cytokine/μg cell protein) comparison of Betamethasone, KNa and S-KA are expressed in Table S4.

**Fig 3.**
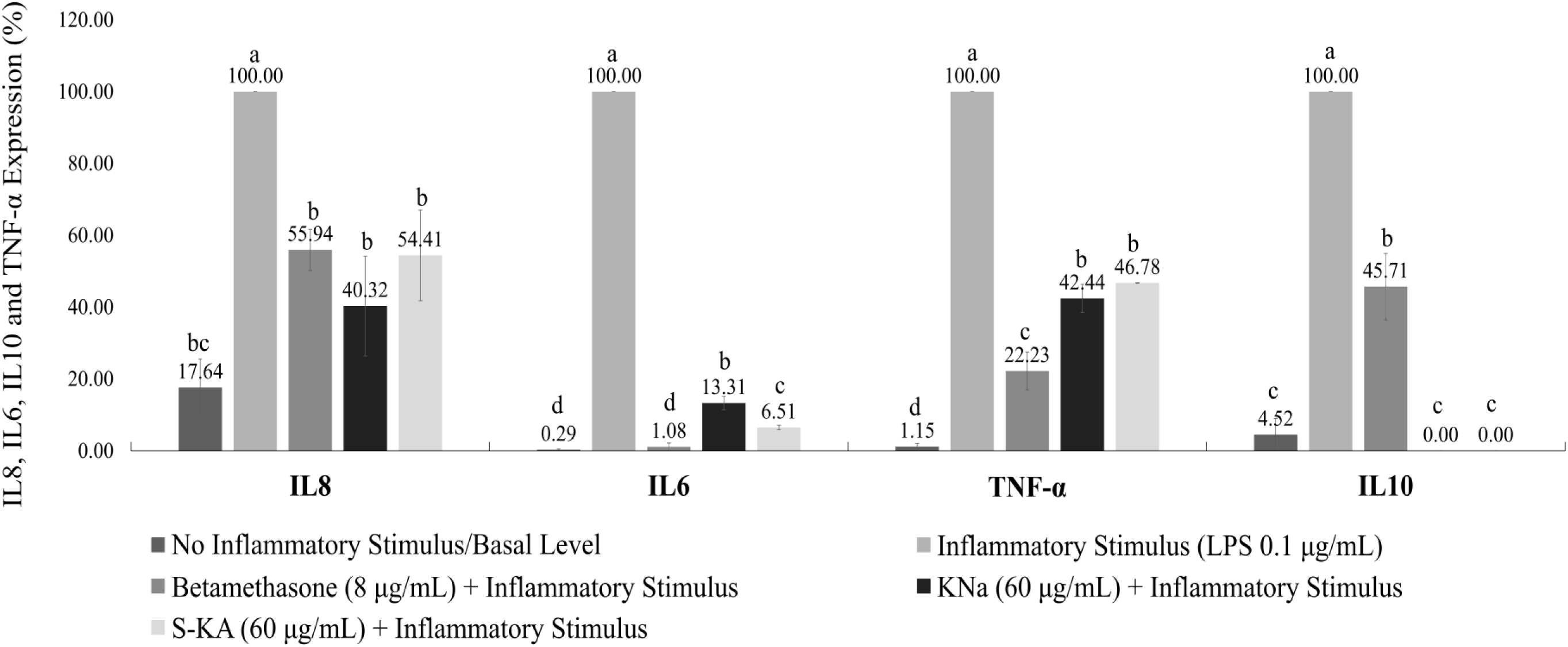
Effect of Kaurenoic Acid (S-KA) and Kaurenoic Sodium Salt (KNa) on IL8, IL6, IL10 and TNF-α Macrophages Expression (%). Different letters (a, b, c, d) for statistically significant differences (p < 0.05).

When responding to pro-inflammatory stimuli, IL-8 can be expressed by macrophages. In fact, as can be observed in Fig 3, the IL-8 expression between cells exposed to an inflammatory stimulus (LPS, considered as 100% of IL-8 expression), compared to the expression on basal level (17.64 ± 7.91%), revealed significant differences (*p* < 0.05). Nevertheless, no statistical differences were observed between cells without inflammatory stimulus or S-KA (54.41 ± 12.62% expression values), and KNa (40.32 ± 13.93% expression values) at 60 μg/mL of S-KA and KNa. The treatment with 8 μg/mL of betamethasone (used as positive control) led to a decrease in IL-8 expression to 55.94 ± 5.72%. Concerning cytokine inhibition (Supplementary Table S4), results suggest that betamethasone impact on IL-8 is close to the one obtained for KNa and S-KA (44.06 ± 5.72, 59.68 ± 13.93 and 45.59 ± 12.62% inhibition, respectively). The corresponding values for IL-8 quantification were 610.03 ± 38.39, 457.73 ± 166.78 and 612.11 ± 172.05 pg IL-8/μg cell protein for betamethasone, KNa and S-KA, respectively.

For IL-6 studies, betamethasone inhibited 98.71 ± 0.86% its expression (Supplementary Table S4). Although both KNa and S-KA also decreased the IL-6 expression significantly, the values did not reach basal levels (Fig 3). Comparing S-KA with KNa inhibition values (Table S4), S-KA inhibited IL-6 expression more effectively, namely 93.49 ± 0.67 and 86.69 ± 1.89% inhibition (quantified as 3.32 ± 0.63 and 6.87 ± 1.60 pg IL-6/μg cell protein) for S-KA and KNa, respectively. Human TNF-α is a cytokine which can exert regulatory and cytotoxic effects.

TNF-α expression values (Fig 3) under LPS on macrophages decreased with betamethasone, KNa and S-KA exposure from 100% of expression to 22.23 ± 5.23, 42.44 ± 3.91 and 46.78 ± 0.10% (quantified as 18.45 ± 5.93, 34.68 ± 6.40 and 37.95 ± 3.62 pg TNF-α /μg cell protein), respectively (Table S4). Considering these expression results, no significant differences occurred among KNa and S- KA. Betamethasone decreased inflammation values, but not to basal levels of TNF- α, as can be seen at Fig 3.

Previous *in vitro* anti-inflammatory studies of kaurenoic acid at 25 µM from *Copaifera* spp. oleoresin on macrophages showed the inhibition of IL-6 (inhibition value of 11.2 ± 5.7%) and the increase of IL-10 production (Vargas et al., 2015). The functions of IL-10 include inhibition of macrophage-mediated cytokine synthesis, which can predict its anti-inflammatory effect. Contrarily, our results (Table S4) suggest a total IL-10 inhibition (100%) by both the kaurenoic acid and its sodium salt, and the corticosteroid used as a positive control (betamethasone) resulted in 54.29 ± 9.26% of IL-10 inhibition.

The *in vivo* anti-inflammatory activity of kaurenoic acid (1–10 mg/kg) extracted from *Sphagneticola trilobata* L. is confirmed by its ability to reduce cytokine (TNF-α, IL-1β and IL-33) release in mice after lipopolysaccharide-induced peritonitis. Interleukin-10 levels were also evaluated by Borghi *et al*. and it was observed an increase in its expression (Borghi et al., 2021). Other research based on oral administration of KA-enriched *Annona tomentosa* extracts from Amazon (100 mg/kg), also evidenced the reduction of TNF-α and IL-1β by 66 % and 35 % inhibition, respectively, on dermal inflammation mice models (Dalenogare et al., 2019).

In summary, the results abovementioned demonstrate that the anti-inflammatory activity of both kaurenoic acid and its sodium salt on interleukins IL-8 and IL-6 was similar to the inhibitory values observed by betamethasone. However, when compared to betamethasone, differences in the anti-inflammatory performance of kaurenoic acid and salt on TNF-α and IL-10 were observed, as both acid and salt inhibited at a lower extent TNF-α and completely inhibited IL- 10 comparatively to betamethasone. Hence, future research works may focus on understanding from a mechanistic point of view, the differences in the inflammatory hallmark between KA and betamethasone, to understand whether the molecular pathways are the same.

#### 3.5.3. Antimicrobial activity

The lowest minimum inhibitory concentration (MIC) for S-KA was observed for *S. aureus* and *S. epidermidis* at 25 μg/mL (Table 3). For KNa the MIC for *S*. *aureus* was 50 μg/mL and 25 μg/mL for *S*. *epidermidis.* Concerning the other microorganisms (*E. coli*, *P. aeruginosa* and *C. albicans*), no antimicrobial effect was observed for the concentrations tested.

**Table 3.**
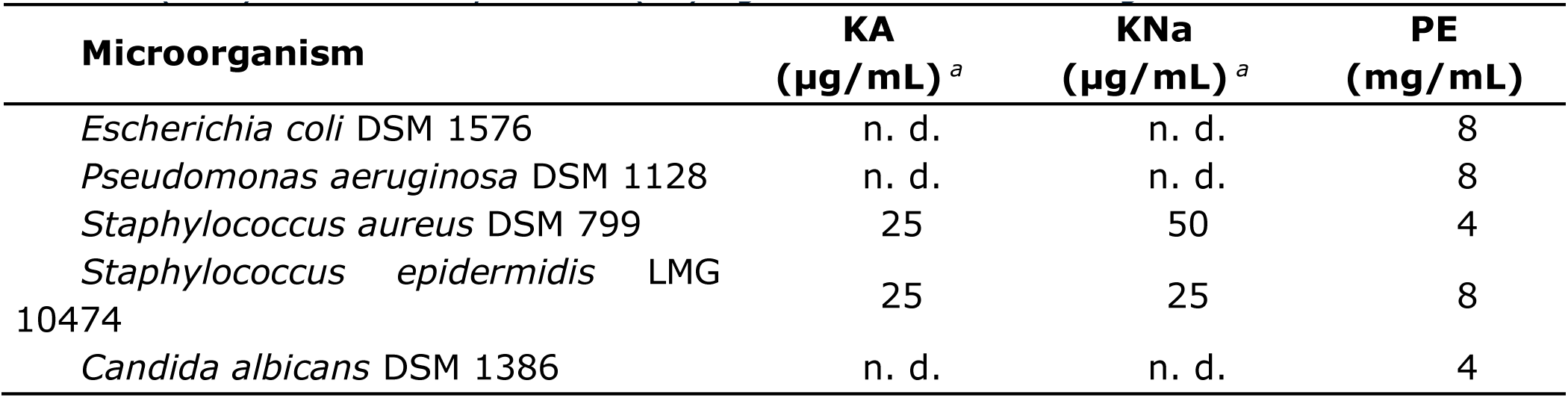
Minimum inhibitory concentration (MIC) for Kaurenoic acid (S-KA), Karenoic salt (KNa) and Phenoxyethanol (PE) against several microorganisms.

Antimicrobial activity of kaurenoic acid (KA) against *S*. *aureus* and *S*. *epidermidis* has already been demonstrated and previously described against Gram-positive organisms such as *S. aureus, S. epidermidis,* and *Bacillus subtilis* when assaying this compound after isolation from leaves of *Smallanthus sonchifolius* and purification by TLC (Padla et al., 2012). These authors using the disk diffusion method, observed MIC values of 125, 250 and 1000 μg/mL respectively for *S. aureus, S. epidermidis,* and *B. subtilis*. Furthermore, when they tested *C. albicans*, it was not also found antimicrobial activity.

Additionally, other authors (De Andrade et al., 2011; Moreira et al., 2016) also determined antimicrobial activity of KA against oral pathogens, mainly species of the genera *Streptococcus*. These authors used the microdilution broth method and determined MIC values of 10 μg/mL for several Streptococci, 100 μg/mL for *Streptococcus salivarius* and 200 μg/mL for *Enterococcus faecalis*.

## 4. Conclusions

In conclusion, this research study confirmed that kaurenoic acid can be obtained from synthetic biology (S-KA) with high-purity levels (purity obtained by GC-MS was 99.06 ± 0.01%). The natural origin of the commercial KA (G-KA) explains the presence of small additional peaks in its GC-MS analysis and, consequently a purity of 98.83 ± 0.04%. Moreover, to overcome the low water solubility of kaurenoic acid, its sodium salt was synthesized (KNa) with a purity value of 99.88 ± 0.01%. The biological activities proved the bioactive potential of kaurenoic acid as an anti-inflammatory and antimicrobial compound. Both kaurenoic acid and its sodium salt at 60 μg/mL decreased interleukins IL-8 and IL-6 and prevented *S. aureus* and *S. epidermidis* growth at lower concentrations (25 µg/mL). Although the kaurenoic acid sodium salt showed higher solubility in water, it did not show greater bioactivity.

## Funding

This research was funded by the European Regional Development Fund (ERDF), through the Operational Program for Competitiveness and Internationalization (COMPETE 2020) and Portugal 2020, under the Alchemy Project (POCI-01-0247-FEDER-027578).

## CRediT authorship contribution statement

**Lígia L. Pimentel**: Conceptualization, Methodology, Data curation, Investigation, Supervision, Writing – original draft, Writing – review & editing. **Francisca S. Teixeira**: Investigation, Writing – original draft, Writing – review & editing. **Ana M. S. Soares**: Methodology, Data curation, Investigation, Writing – original draft, Writing – review & editing. **Paula T. Costa**: Investigation, Writing – original draft, Writing – review & editing. **Ana Luiza Fontes**: Investigation, Writing – original draft, Writing – review & editing. **Susana S. M. P. Vidigal**: Investigation, Writing – original draft, Writing – review & editing. **Manuela E. Pintado**: Funding acquisition. **Luis M. Rodríguez-Alcalá**: Conceptualization, Methodology, Formal analysis, Data curation, Writing – original draft, Writing – review & editing, Supervision.

## Declaration of competing interest

The authors declare no conflict of interest. The funders had no role in the design of the study; in the collection, analyses, or interpretation of data; in the writing of the manuscript; or in the decision to publish the results.

## Data availability

Data will be made available on request.

## Abbreviations

KA: Kaurenoic Acid
S-KA: Synthetic Kaurenoic Acid
G-KA: Commercial Kaurenoic Acid (Natural Extract)
KNa: Kaurenoic Sodium Salt
QSAR: Quantitative Structure-Activity Relationship
IL: Interleukin
TNF-α: Tumor Necrosis Factor Alpha
LPS: Lipopolysaccharides
DMSO: Dimethyl Sulfoxide
EtOH: Ethanol
EtOAc: Ethyl Acetate
AcO: Acetone
Hxn: Hexane
DCM: Dichloromethane
FTIR-ATR: Fourier Transform Infrared Spectroscopy-Attenuated Total Reflectance
DSC: Differential Scanning Calorimetry
XRD: X-ray Diffraction
LC-ESI-QTOF: Liquid Chromatography Electrospray Ionization Quadrupole Time-of-Flight
GC-MS: Gas Chromatography-Mass Spectrometry
HRMS: High Resolution-Mass Spectrometry
RPMI: Roswell Park Memorial Institute (used for cell culture medium)
FBS: Fetal Bovine Serum
DMEM: Dulbecco’s Modified Eagle Medium
HaCaT: Human Keratinocyte Cell Line

## Suplementary information

**Fig S1.**
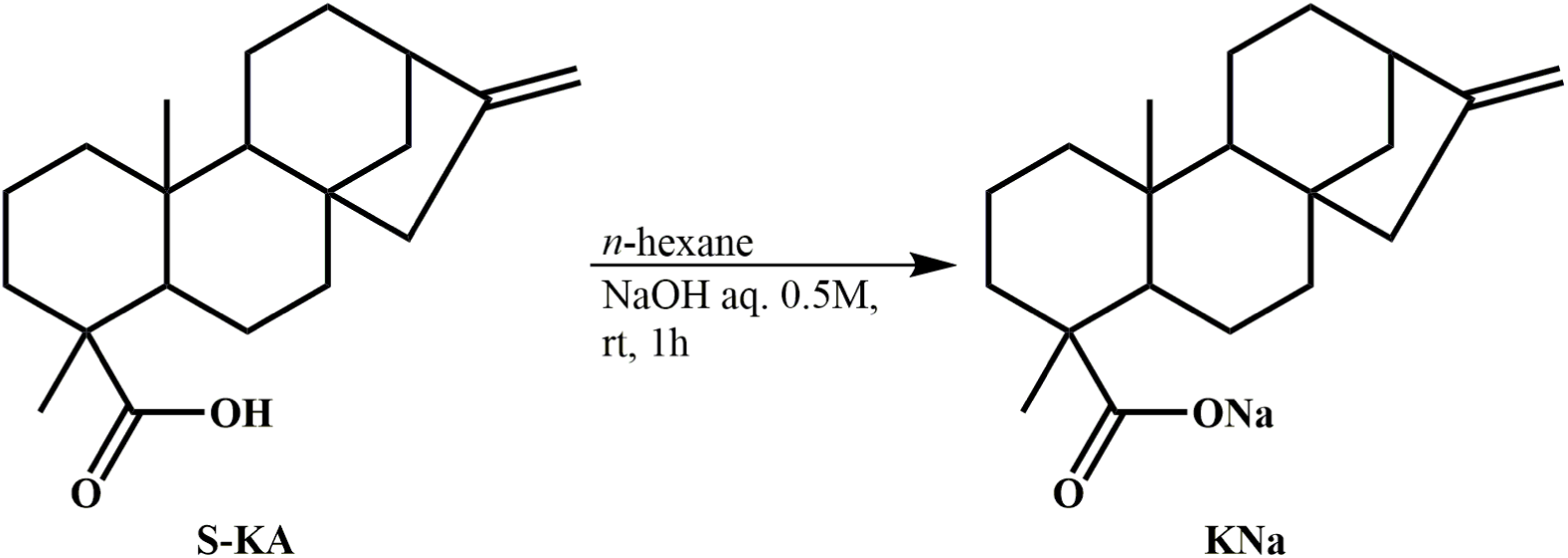
Synthesis conditions for Kaurenoic Sodium Salt (KNa).

**Fig S2.**
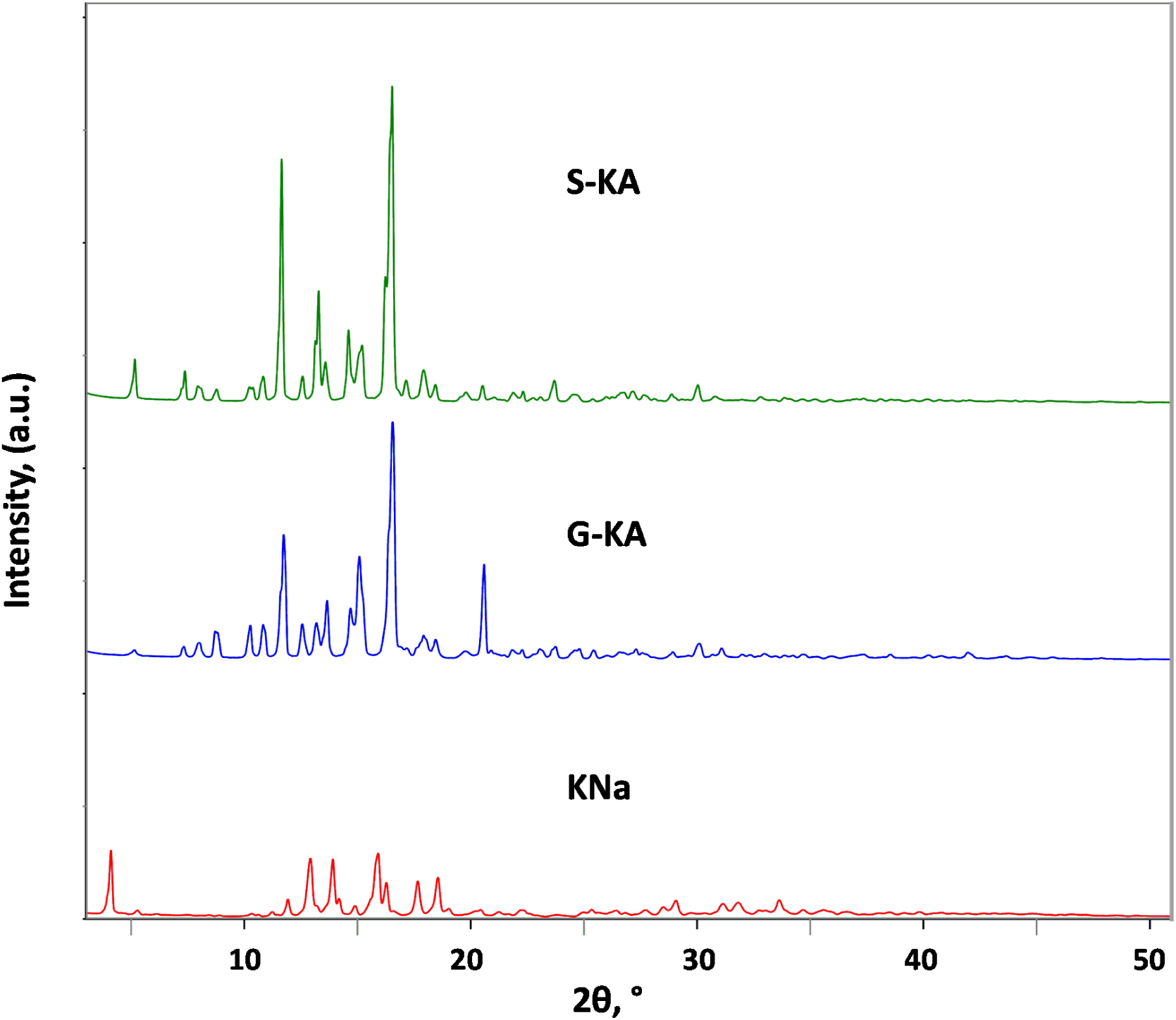
X-Ray diffraction patterns of S-KA, G-KA and KNa.

**Fig S3.**
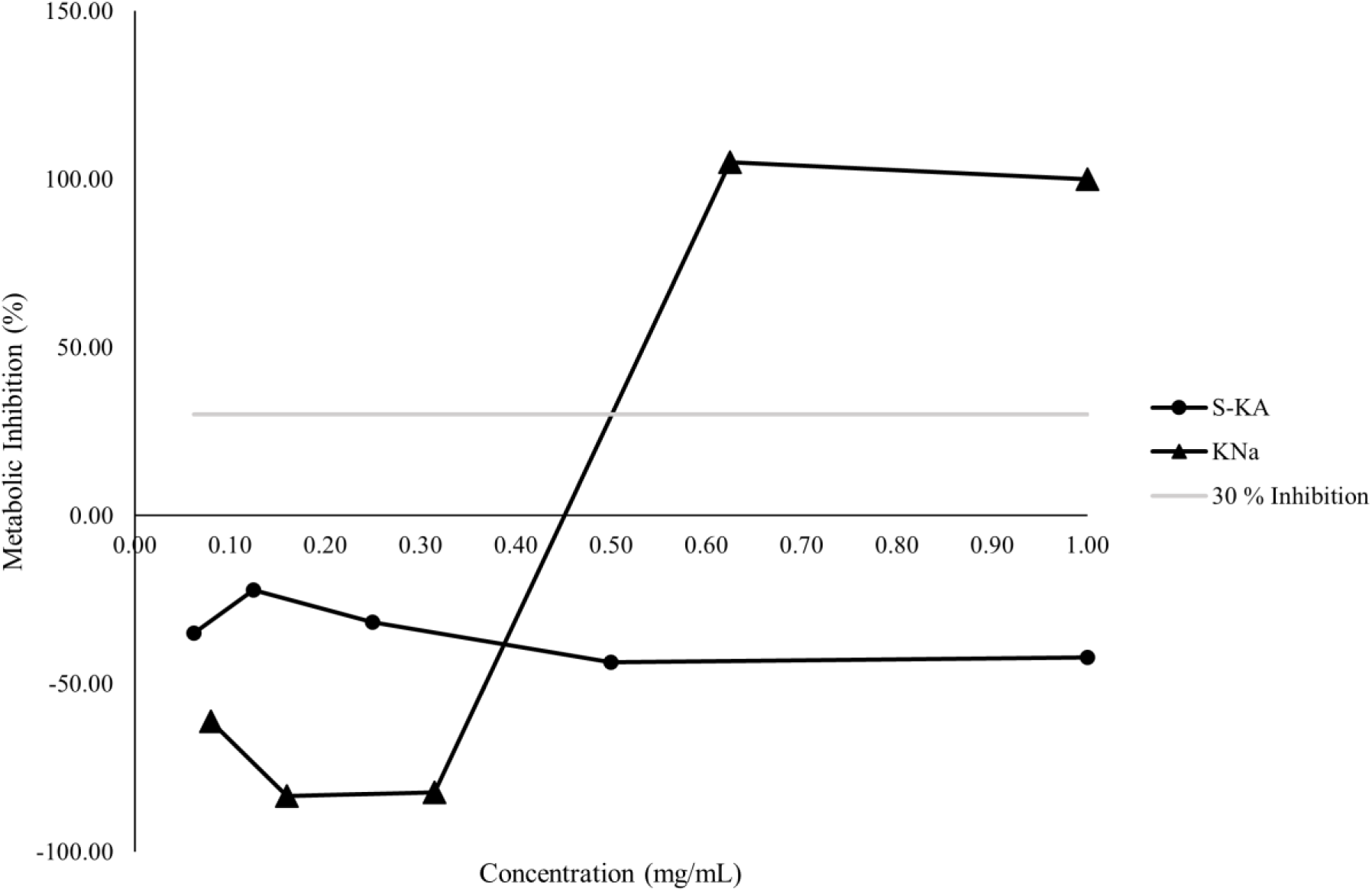
Metabolic Inhibition (%) of Kaurenoic Sodium Salt (KNa) and Kaurenoic Acid (S-KA) on Macrophages.

**Fig S4.**
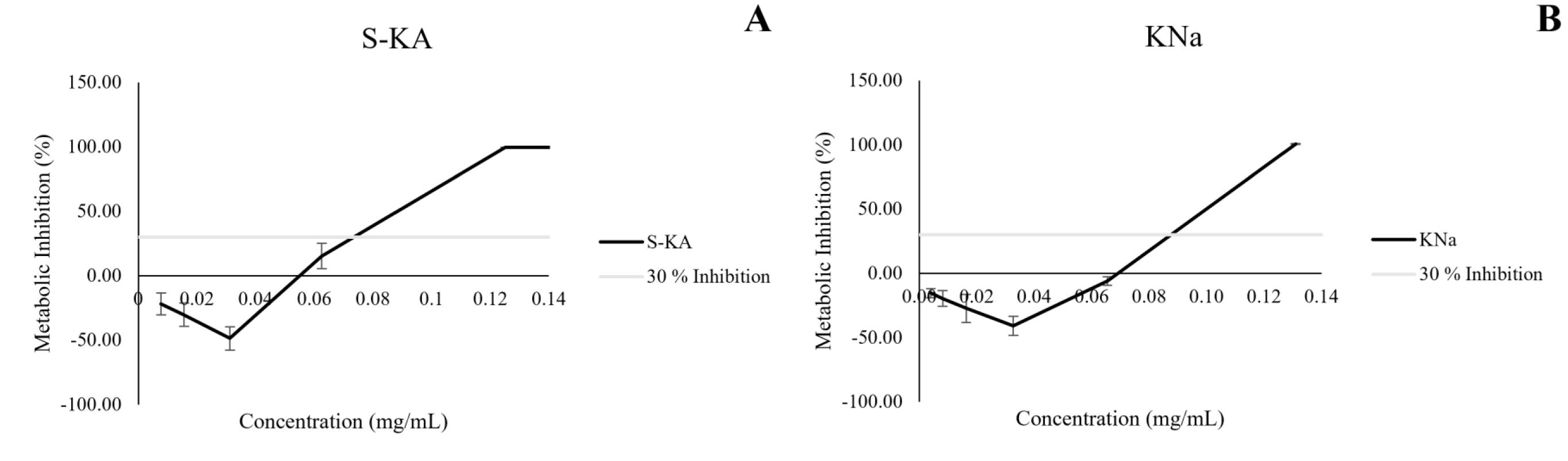
Metabolic Inhibition (%) of A. Kaurenoic Acid (S-KA) and B. Kaurenoic Salt (KNa) on Keratinocytes.

**Table S1.**
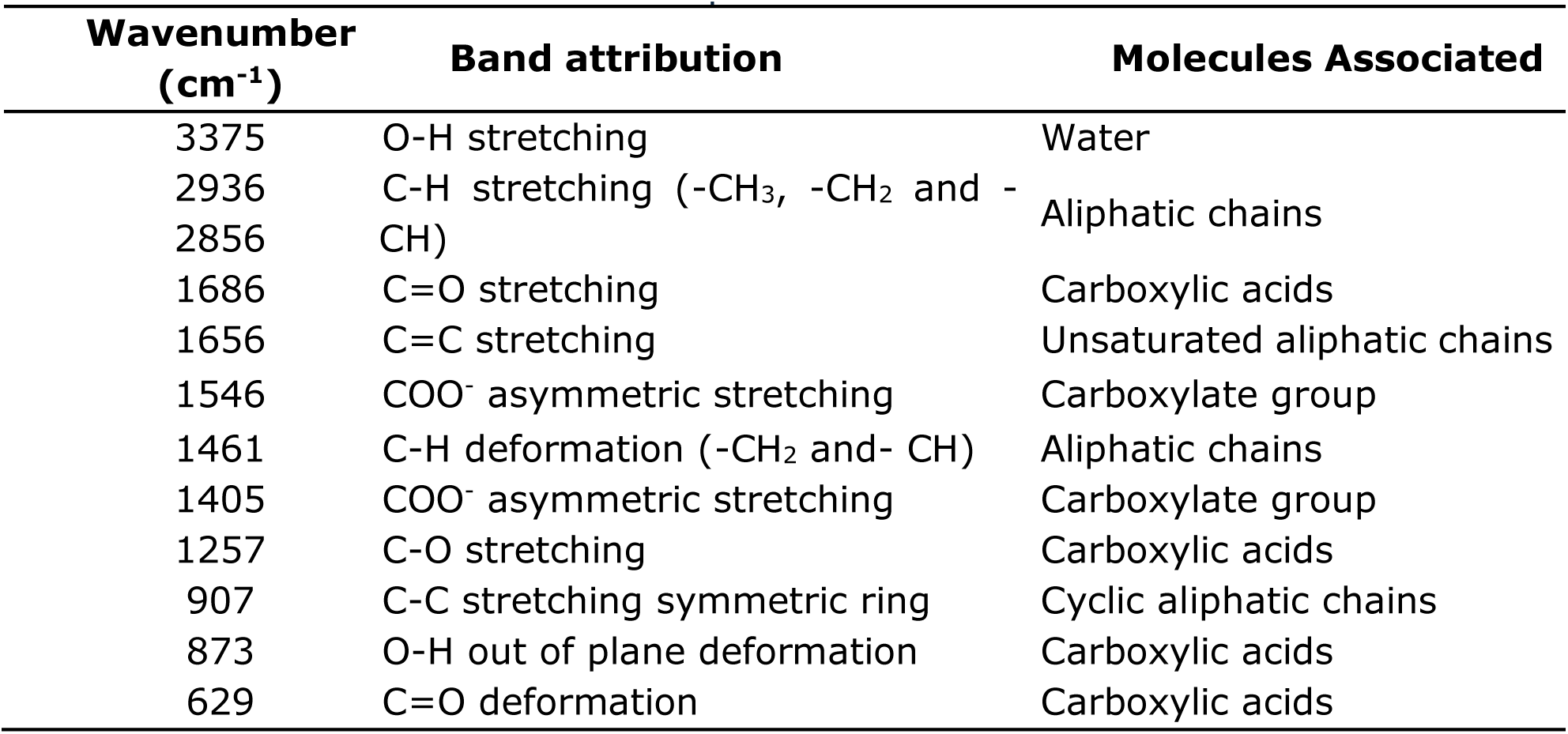
FTIR-ATR vibrational bands interpretation.

**Table S2.**
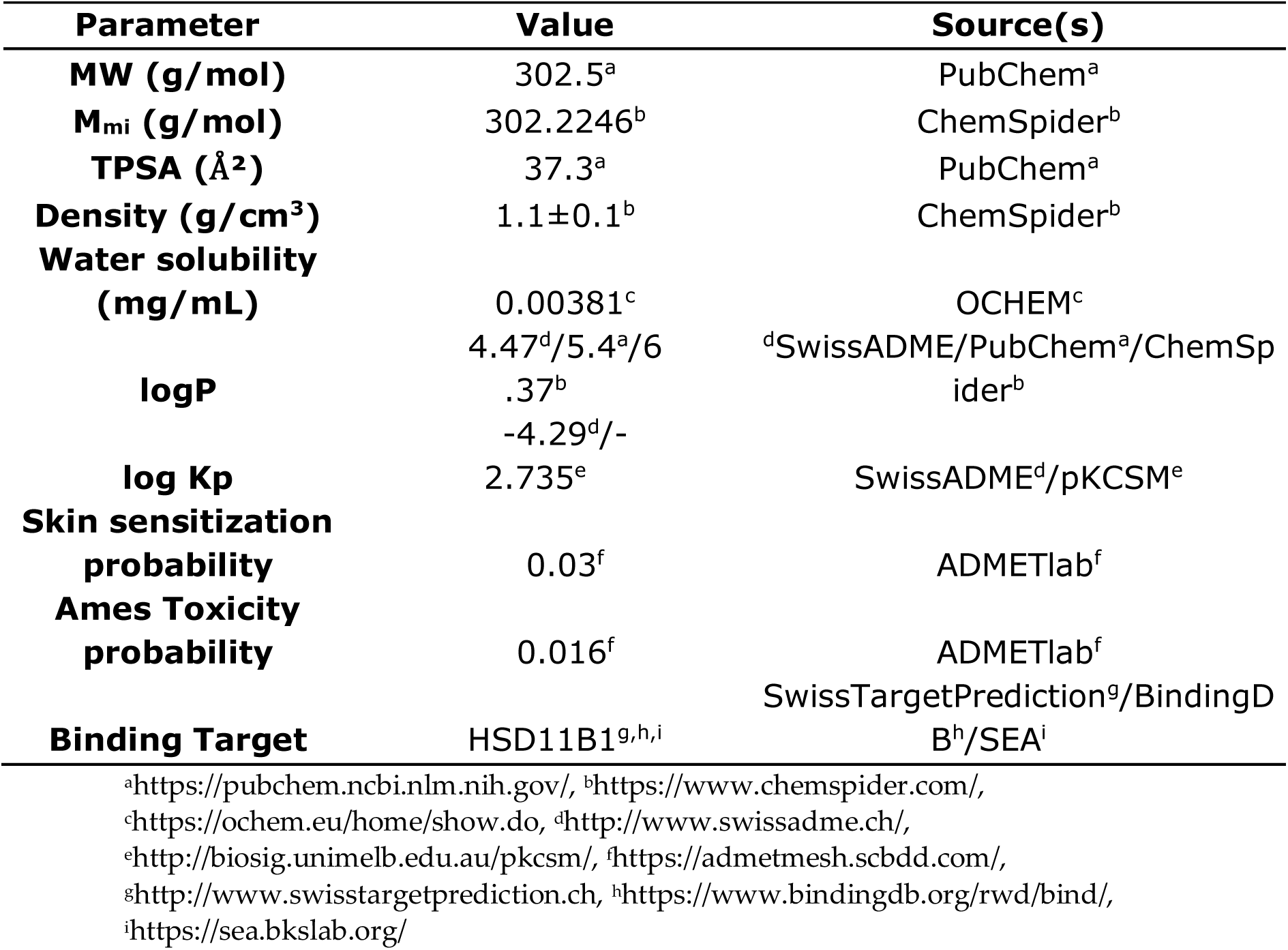
Physicochemical properties and quantitative structure-activity relationship (QSAR) of Kaurenoic Acid.

**Table S3.**
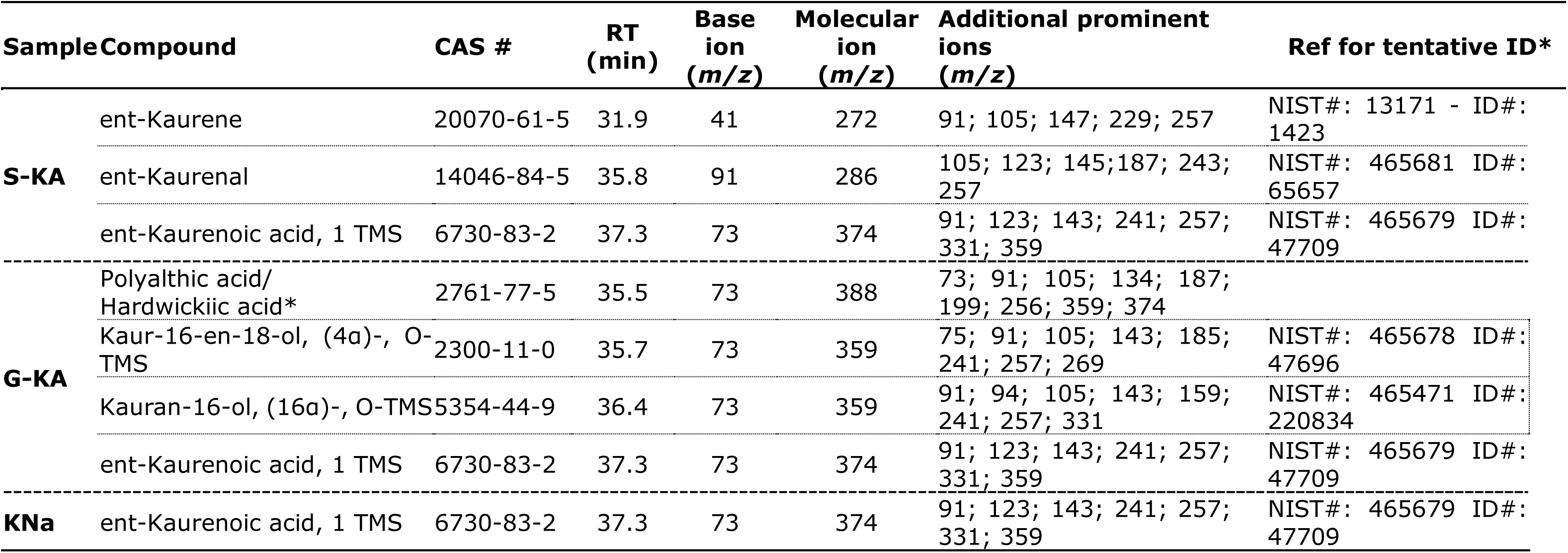
Retention time and mass spectra data for S-Ka, G-KA and KNa compounds identified by GC-MS. *tentative identification; no reference spectra available

**Table S4.** Cytokine inhibitory activity (%) and Cytokine Concentration (pg cytokine/μg cell protein) comparison of Betamethasone, KNa <u>and S-KA.</u>.

|  |  | **IL8** |  |  | **IL6** |  |  | **TNF-α** |  |  | **IL10** |  |  |
| --- | --- | --- | --- | --- | --- | --- | --- | --- | --- | --- | --- | --- | --- |
| Cytokine<br>Inhibitory Activity<br>(%) | Betamethasone (8 μg/mL) +<br>LPS | 44.06 | ± | 5.72 | 98.71 | ± | 0.86 | 77.77 | ± | 5.23 | 54.29 | ± | 9.26 |
|  | KNa (60 μg/mL) + LPS | 59.68 | ± | 13.93 | 86.69 | ± | 1.89 | 57.56 | ± | 3.91 | 100* |  |  |
|  | S-KA (60 μg/mL) + LPS | 45.59 | ± | 12.62 | 93.49 | ± | 0.67 | 53.22 | ± | 0.10 | 100* |  |  |
| Cytokine<br>Concentration<br>(pg cytokine/μg<br>cell protein) | Betamethasone (8 μg/mL) +<br>LPS | 610.03 | ± | 38.39 | 1.07 | ± | 0.07 | 18.45 | ± | 5.93 | 0.20 | ± | 0.02 |
|  | KNa (60 μg/mL) + LPS | 457.73 | ± | 166.78 | 6.87 | ± | 1.60 | 34.68 | ± | 6.40 | 0 |  |  |
|  | S-KA (60 μg/mL) + LPS | 612.11 | ± | 172.05 | 3.32 | ± | 0.63 | 37.95 | ± | 3.62 | 0 |  |  |

\* Complete inhibition in all replicates

## Notes

### Competing Interest Statement

The authors have declared no competing interest.

